# NeoOmni Atlas: an integrated database for pan-cancer neoantigen derived from proteogenomics analysis

**DOI:** 10.64898/2026.10.09.757967

**Authors:** Tao Zheng, Zhi Ran, Jing Zeng, Tao Wang, Shiyang Fu, Lihua Cai, Dinghong Wu, Jiaxiang Ding, Tang Liu, Zhihong Li, Kai Yuan, Shaofeng Lin, Haodong Xu

## Abstract

Tumor neoantigens derived from diverse molecular aberrations are important targets for personalized cancer immunotherapy, but existing resources incompletely integrate mass-spectrometry-supported canonical and non-canonical antigen evidence across cancer types. NeoOmni Atlas is an interactive pan-cancer T-cell epitope resource, built via proteogenomic reanalysis of 7,561 immunopeptidomics datasets comprising 511 million raw spectra from 2,341 samples across 26 cancer types. The analysis yielded 872,414 eluted canonical peptides and 198,961 non-canonical peptides spanning 18 molecular alteration classes, along with 759,386 unique ligands presented by 322 human leukocyte antigen (HLA) alleles. Relative to existing repositories, NeoOmni Atlas covers a broader range of cancer types, includes 60 additional HLA alleles, and catalogs 369,781 novel ligands. A computational tumor-specific filtering framework identified 316,126 tumor-associated epitopes and 61,231 tumor-specific epitopes. Tumor-specific epitopes showed stronger predicted binding affinity than tumor-associated epitopes for HLA class I A and B alleles, and 65 conserved tumor-specific epitopes recurrent across more than 10 tumor types represent potential shared antigen candidates. NeoOmni Atlas also prioritizes candidate tumor-associated antigens including *PMEL* (gp100:209–217) and *CSPG4*, and supports browsing, searching, and downloading by cancer type, human leukocyte antigen allele, peptide sequence, and source event. NeoOmni Atlas is freely accessible at http://modinfor.com/NeoOmni-Atlas/.

## Introduction

Many antitumor treatments rely on T-cell immunity [1]. Antigen specificity originates from binding between a T-cell receptor (TCR) and a peptide-loaded human leukocyte antigen (HLA) molecule [2,3]. This interaction underlies immune checkpoint inhibitors, adoptive tumor-infiltrating lymphocyte transfer, genetically modified TCR-expressing T cells, as well as chimeric antigen receptor (CAR) T cells [2,4–7]. Neoantigens derived from somatic mutations, transcriptional dysregulation, or non-canonical translation may show tumor enrichment and reduced central tolerance, making them attractive candidate targets for precision immunotherapy [8,9]. Early trials of individualized neoantigen vaccines in melanoma, glioblastoma, non-small-cell lung cancer, and pancreatic ductal adenocarcinoma have reported favorable safety and initial efficacy [10–14], reinforcing the need for robust HLA-informed antigen resources.

Neoantigen discovery faces limitations. In silico pipelines based on variant calls, transcriptomics, and binding prediction remain prone to false positives [15–18]. In immunopeptidomics, chromatographic peptide separation precedes tandem mass-spectrometric analysis (LC-MS/MS). This workflow detects HLA-presented peptides directly, confirms predicted antigens, and uncovers cryptic or non-canonical peptides [15]. Large-scale MS/MS datasets may contain underutilized peptide-level information: mass-tolerant searches, spectral library analyses, expanded search spaces, and standardized reanalysis workflows can recover additional peptide evidence from previously unidentified spectra, although the extent is dataset- and workflow-dependent [15,19–22]. This issue is particularly relevant in immunopeptidomics, because HLA-presented ligands can arise not only from annotated proteins but also from variants, junctions, alternative open reading frames (ORFs), RNA-level events, and pathogen-associated sequences. Existing resources, including HLA Ligand Atlas, SysteMHC Atlas, and caAtlas [22–24], have advanced canonical HLA ligand annotation, while IEAtlas, MicroEpitope, FusionNeoAntigen, and TSNAdb cover selected non-canonical sources [25–28]. However, these resources differ in data origin, search strategy, event coverage, and direct MS support for the reported peptides. A unified immunopeptidomics-supported pan-cancer resource that harmonizes ligandome data, reprocesses spectra, and annotates canonical and non-canonical antigen sources remains needed.

Here, we present NeoOmni Atlas, an interactive pan-cancer tumor (neo)antigen resource available at http://modinfor.com/NeoOmni-Atlas/. By reanalyzing over 7,500 immunopeptidomics mass spectrometry raw files comprising 511 million raw spectra from 2,341 samples across 26 cancer types, we identified 1,071,375 MS-eluted peptides, including 872,414 canonical and 198,961 non-canonical peptides, and assigned 759,386 unique HLA-presented peptides to 322 HLA alleles. Using a computational tumor-specific filtering framework, we identified 61,231 tumor-specific epitopes (TSEs; non-canonical/tumor-restricted) and 316,126 tumor-associated epitopes (TAEs; canonical/tumor-exclusive). NeoOmni Atlas further documents 369,781 novel HLA ligands relative to existing repositories, incorporates 60 additional HLA alleles, and provides peptide-, HLA-, cancer-type-, event-, and post-translational modification (PTM)-level annotations for browsing, searching, and downloading. By combining direct MS evidence with source-event annotation and predicted HLA association, the resource is intended to support systematic hypothesis generation for tumor antigen discovery. These annotations support antigen prioritization rather than experimental validation of clinical tumor specificity.

## Database construction and methods

### Sources of Immunopeptidomics Datasets

We searched jPOST, PRIDE, PeptideAtlas, MassIVE, and iProX [29–33] using keywords including “HLA”, “MHC”, “immunopeptidomics”, “HLA ligand”, and “HLA epitopes”. Datasets were manually curated and retained only if they reported MS-based identification of HLA-bound peptides, typically after immunoaffinity purification with anti-HLA antibodies. In total, 117 immunopeptidomics datasets were collected, and the full list is available on the NeoOmni Atlas Download page.

### Sample Information Collection

For each dataset, sample-level metadata were manually extracted from the associated publications, including cancer or tissue type, HLA class I/II alleles, sample identifiers, and PubMed ID. Only samples with clearly reported HLA genotype and tissue origin were retained. When multiple preparations, instruments, or conditions were available for the same patient or cell line, they were retained as separate sample-level entries for downstream processing. A detailed summary is provided in Supplementary Table S1. These curated metadata fields were subsequently used to constrain HLA deconvolution, assign cancer-type labels, and support cancer-specific browsing in the web interface.

### Sources of 18 Neoantigen-Generating Non-canonical Events

To characterize non-canonical antigen diversity, we integrated 18 non-canonical/additional antigen source classes from public resources and literature-derived datasets. These source classes include somatic alterations from The Cancer Genome Atlas (TCGA)/Xena [34]; gene fusions and transposable element fusions [35,36]; lncRNA-, pseudogene-, and alternative ORF-related events from nuORFdb and literature-supported overlapping translation analyses [37,38]; RNA editing events from RADAR [39]; and pathogen- or virus-derived sequences from MicroEpitope [25]. These sequences were derived from multiple public resources rather than fully matched sequencing data from the same immunopeptidomics samples, reflecting the heterogeneous origin of the integrated non-canonical source classes. Event sources, event numbers, and construction rules are summarized in Supplementary Table S2, enabling transparent interpretation of the expanded non-canonical search space.

### Event-specific construction of canonical and non-canonical search spaces

Canonical peptides were defined as peptides mapped to annotated protein-coding sequences in the reference proteome and assigned to the Canonical_ORF category. Non-canonical peptides were identified from expanded search spaces derived from the 18 non-canonical/additional source classes. Search-space construction was event-specific: local variant, intronic, breakpoint, or junction templates considered strand-aware and six-frame translation where applicable; annotated ORF classes used source database coordinates and strand information; and source-provided peptide or protein entries were incorporated according to available annotation. Mutation- and junction-derived templates were constructed to preserve the altered residue or junction-spanning sequence where applicable, and peptide source annotations were assigned after database searching rather than solely from event labels. Detailed rules are provided in Supplementary Table S2 and Supplementary Figure S1. This event-specific design was used to avoid treating all non-canonical events as a single homogeneous ORF set and to clarify how variant-, junction-, intronic-, pseudogene-, pathogen-, and alternative ORF-derived templates contributed to the final searchable database.

### Definition of epitope categories

TAEs were operationally defined as canonical tumor-derived peptides not detected in the compiled normal/benign peptide reference set. Candidate TAEs were derived from peptides mapped to canonical protein-coding regions in tumor samples and filtered against the compiled normal/benign peptide background; therefore, “tumor-exclusive” refers to absence from our compiled normal reference peptides rather than proof of universal absence from all possible normal tissues. TSEs were defined as tumor-type-specific and tissue-context-dependent non-canonical peptides after removal of peptides detected in corresponding normal/benign references and the complete TAE peptide set. Normal-Epitope Sources (NES) include normal-only peptides and peptides co-expressed in malignant and benign contexts. TAE, TSE, and NES were treated as mutually exclusive final annotation categories. This classification was used consistently in downstream analyses and figure legends to avoid mixing canonical tumor-exclusive candidates, non-canonical tumor-context-restricted candidates, and normal or co-expressed epitope sources.

### Epitope Identification

Peptide identification was performed using pFind/open-pFind [40] with a unified workflow across datasets to reduce search-engine-dependent variability. For low-resolution datasets, precursor-ion and fragment-ion mass tolerances were 5 ppm and 0.5 Da, respectively; high-resolution datasets used corresponding tolerances of 20 ppm and 0.05 Da. High-resolution datasets were additionally analyzed in open-search mode to identify peptides bearing post-translational modifications. HLA-I and HLA-II peptides were limited to 8–15 and 8–25 residues, respectively, consistent with typical ligand-length distributions.

### Epitope identification quality control

Peptide identification was performed at the sample level. Samples from the same patient but processed differently or analyzed on different instruments were searched separately. We applied a false discovery rate (FDR) cutoff below 1% to each identification level: peptide–spectrum matches, peptides, and proteins. Following integration, spectra were ranked again according to Q-values, the combined FDR was recalculated, and spectra that failed the cutoff were removed. This integrated quality-control strategy was used to maintain a consistent confidence level across heterogeneous public datasets. The pFind-based workflow was further evaluated using cross-engine comparisons, overlap with established immunopeptidomics resources, and retention-time consistency analyses, providing additional support for the reliability of peptide identification.

### Assignment of Epitopes to Specific HLA Alleles

Because most samples were multi-allelic, we used NetMHCpan 4.1 [16] and NetMHCIIpan 4.3 [41] for genotype-informed HLA deconvolution. Experimentally identified peptides were evaluated only against HLA alleles expressed by the corresponding sample, and the allele with the lowest predicted %Rank was assigned as the primary predicted presenting allele. Default NetMHCpan thresholds were used for strong, weak, and non-binders: for HLA-I predictions, %Rank values below 0.5 indicated strong binding, whereas values ranging from 0.5 to 2 indicated weak binding. The corresponding thresholds for HLA-II were below 1 and from 1 to 5, respectively. These assignments should be interpreted as prediction-based HLA associations rather than definitive experimental HLA restrictions.

### Identification of Tumor-restricted Epitope Source Genes by Proteogenomics

For canonical TAE source genes, we applied a tumor-restriction composite scoring strategy integrating TAE counts and transcriptomic abundance within each cancer type. For each gene, the TAE-count score was calculated as follows: TAE score = (TAE count of the gene − mean TAE count across genes) / standard deviation of TAE counts across genes. The expression score was calculated similarly: EXPR score = (expression of the gene − the across-gene mean) / the across-gene standard deviation. The final composite score was obtained by summing the two standardized metrics: composite score = TAE score + EXPR score. This score was used to prioritize genes with both frequent HLA presentation and tumor-associated expression. For TSE-associated events, we did not apply the same gene-expression-based composite score because TSEs arise from heterogeneous non-canonical sources for which comparable gene-level expression matrices are not consistently available. Instead, for events with available TCGA genomic or transcriptomic support, we used a complementary event-level prioritization strategy integrating immunopeptidomics detection and recurrence information, based on counts of detected TSEs and TCGA samples carrying the corresponding event. This strategy prioritizes recurrent MS-supported TSE-associated events without assuming that all non-canonical events can be scored using the same gene-expression-based framework. These analyses were used for candidate ranking and visualization rather than to assert functional immunogenicity.

### Data Organization and Web Deployment

After normalization and integration, peptide entries were organized into canonical peptides, non-canonical peptides, TAEs, TSEs, and NES categories. The curated datasets were integrated into the NeoOmni Atlas web platform, which contains Home, Browse, Search, Download, Guide, and Contact sections. Users can browse by cancer type, HLA allele, post-translational modification (Supplementary Table S3), or tissue type; search by peptide, HLA, event, or PTM; and download data by cancer type, HLA allele, or PTM type, or download complete datasets. Each peptide entry is linked to source annotations, sample context, predicted HLA association, binding information, and available MS/MS evidence.

## Database contents and features

### Workflow and Data Overview of NeoOmni Atlas

NeoOmni Atlas was designed as an integrated proteogenomic and immunopeptidomics-supported database for screening epitopes derived from canonical proteins and 18 non-canonical/additional antigen source classes (**Figure 1**A). Compared with existing resources, it expands the coverage of immunopeptidomics-supported antigen source classes and provides integrated annotations across cancer type, HLA allele, peptide source, event type, and epitope category (Figure 1B). Our analysis leveraged 7,561 immunopeptidomics datasets comprising 511 million raw spectra from 2,341 samples across 26 cancer types and 322 HLA alleles. Event-specific templates were constructed, all MS data were reprocessed under FDR-controlled criteria, and genotype-informed binding prediction assigned ligands to specific HLA alleles. In total, NeoOmni Atlas contains 759,386 unique HLA-presented peptides distributed across 163 HLA-I alleles (mean, 2,392 peptides/allele) and 159 HLA-II alleles (mean, 945 peptides/allele), and identifies 316,126 TAEs and 61,231 TSEs. Relative to existing repositories, NeoOmni Atlas further encompasses 14 unique cancer types, 60 additional HLA alleles, and 369,781 novel HLA ligands. Non-canonical source analysis highlighted broad contributions from alternative ORF-, pseudogene-, somatic alteration-, and lncRNA-related events, including nuORF-related sources (24,831 neoepitopes; 35.3%), pseudogenes (18,562; 26.4%), somatic mutations (15,779; 22.4%), and lncRNAs (11,098; 15.8%) (Figure 1C). Because absolute peptide counts can be influenced by event-class-specific search-space size, we further evaluated search-space-normalized peptide yield across non-canonical event classes (Supplementary Figure S2).

**Figure 1.**
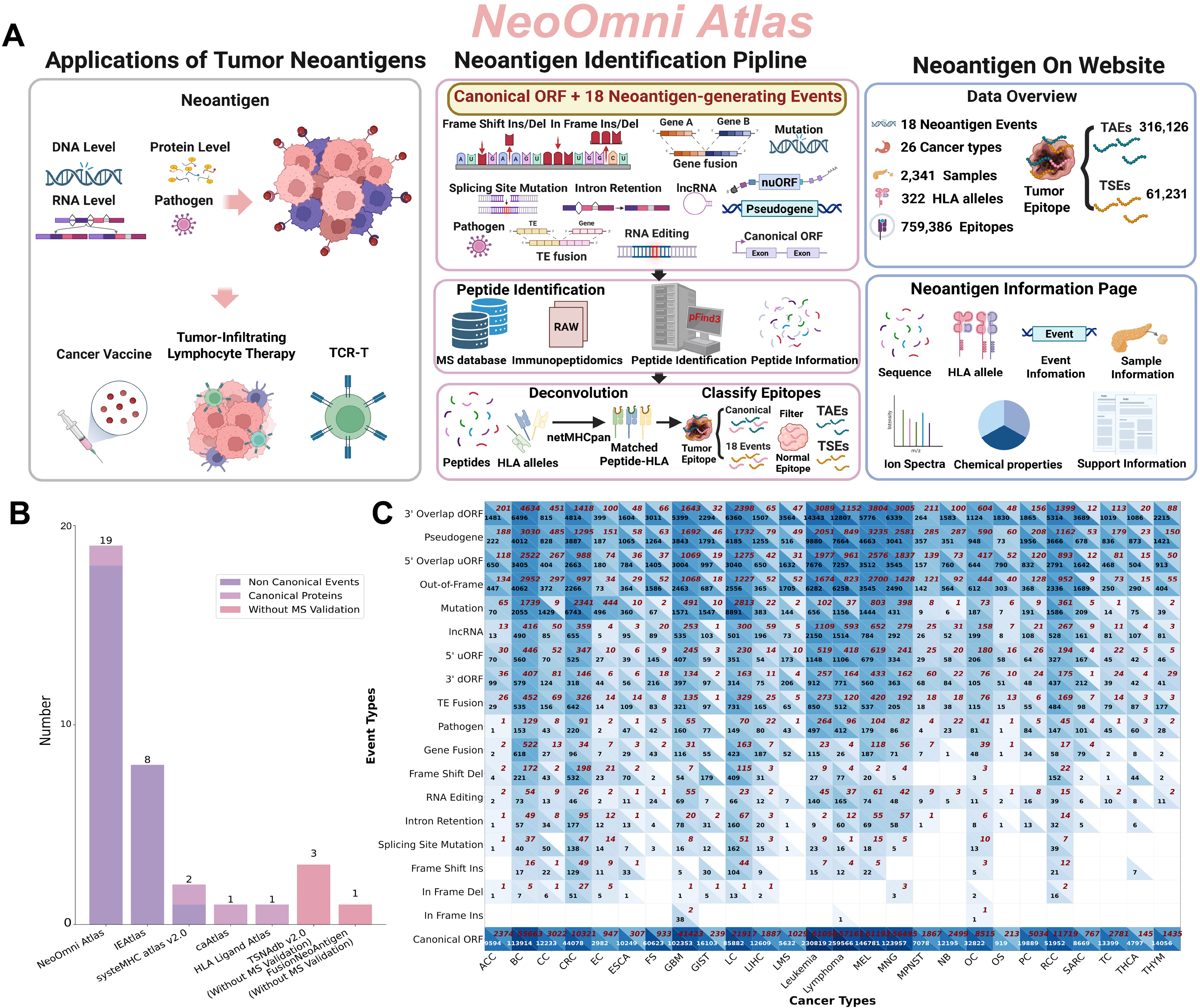
Workflow and data overview of NeoOmni Atlas. (A) Overall workflow of NeoOmni Atlas, including collection of immunopeptidomics datasets, construction of canonical and non-canonical search spaces, peptide identification under FDR-controlled criteria (< 1%), HLA deconvolution, epitope classification, and web deployment. (B) Comparison of the number of neoantigen-generating event types covered by different resources. (C) Cancer-type distribution of TAEs and TSEs generated by different non-canonical event classes. Black numbers indicate the number of TAEs, and red numbers indicate the number of TSEs for each cancer type/event-class combination. Cancer-type abbreviations: ACC, adrenocortical cancer; BC, breast cancer; CC, cervical cancer; CRC, colorectal cancer; EC, endometrial carcinoma; ESCA, esophageal carcinoma; FS, fibrosarcoma; GBM, glioblastoma; GIST, gastrointestinal stromal tumor; LC, lung cancer; LIHC, liver cancer; LMS, leiomyosarcoma; MEL, melanoma; MNG, meningioma; MPNST, malignant peripheral nerve sheath tumor; NB, neuroblastoma; OC, ovarian cancer; OS, osteosarcoma; PC, pancreatic cancer; RCC, renal cell cancer; SARC, sarcoma; TC, testicular cancer; THCA, thyroid cancer; THYM, thymoma. Leukemia and lymphoma are shown as full names. Abbreviations: FDR, false discovery rate; HLA, human leukocyte antigen; TAE, tumor-associated epitope; TSE, tumor-specific epitope; ORF, open reading frame; MS, mass spectrometry; TCR-T, T-cell receptor-engineered T cell; TE, transposable element; lncRNA, long non-coding RNA; nuORF, novel or unannotated open reading frame; uORF, upstream open reading frame; dORF, downstream open reading frame. Alt text: Figure 1 summarizes the NeoOmni Atlas workflow, compares event-type coverage with other resources, and shows cancer-type distributions of TAE and TSE counts across non-canonical event classes.

### Mining Tumor Neoantigen Data Using the NeoOmni Atlas

By separately analyzing immunopeptidomics-validated epitopes in tumor and corresponding normal/benign reference contexts, NeoOmni Atlas identified 316,126 TAEs and 61,231 TSEs. Pan-cancer epitope distributions were heterogeneous. Leukemia exhibited exceptional antigenic richness (> 150,000 total epitopes; **Figure 2**A), suggesting that it may provide a candidate reservoir for adoptive T-cell therapy research. Compared with existing immunopeptidomics repositories, NeoOmni Atlas documented 250,967 newly identified HLA class I ligands and 118,814 HLA class II ligands, totaling 369,781 novel HLA ligands (Figure 2B). This expansion was supported by 37 newly acquired immunopeptidomics datasets, 14 additional cancer types, and 60 additional HLA alleles, including 23 HLA-I and 37 HLA-II alleles (Supplementary Figure S3A–D).

**Figure 2.**
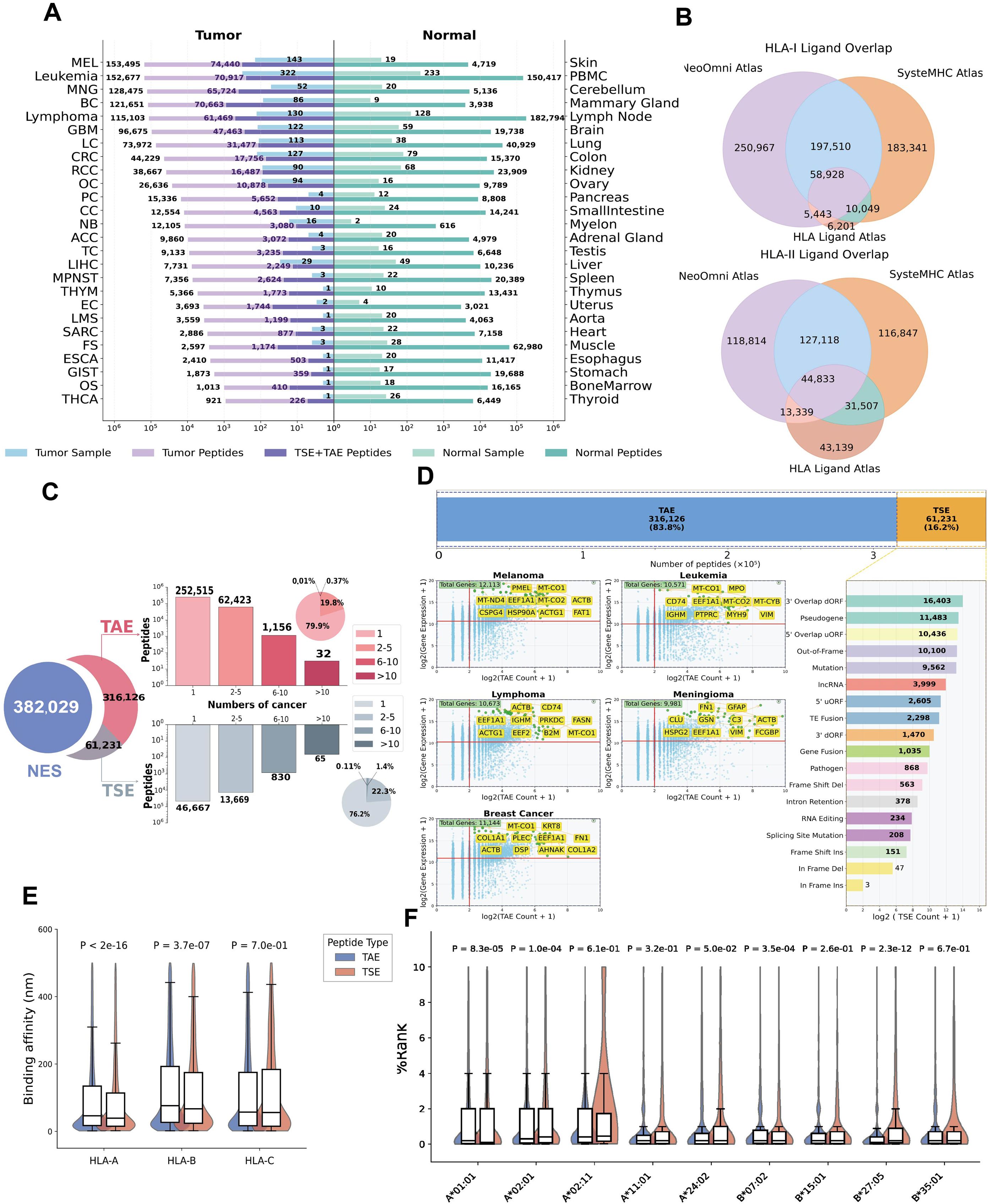
Mining tumor neoantigen epitopes using NeoOmni Atlas. (A) Numbers of TSEs and TAEs identified across tumor samples and corresponding normal/benign references in different cancer types. TAEs were defined as canonical tumor-derived peptides not detected in the compiled normal/benign peptide reference set; TSEs were defined as tumor-type-specific and tissue-context-dependent non-canonical peptides after removal of peptides detected in corresponding normal/benign references and the complete TAE peptide set. Peptide identification was performed under FDR-controlled criteria (< 1%). (B) Overlap of HLA class I and class II epitopes between NeoOmni Atlas and public immunopeptidomics resources, including SysteMHC Atlas and HLA Ligand Atlas. (C) Distribution of TAE, TSE, and NES categories across cancer types. (D) Proportion of TAEs and TSEs, distribution of TSEs across event classes, and proteogenomic composite scoring in representative TAE-enriched cancer types. (E) Comparison of predicted binding affinities of HLA-A, HLA-B, and HLA-C for TSEs and TAEs. Predicted binders were defined using an affinity cutoff of < 500 nM where applicable. (F) Immunogenicity ranking of nine HLA alleles for TSEs and TAEs based on PRIME-HLA-I-predicted peptide-HLA presentation/immunogenic potential using %Rank_bestAllele as the ranking metric; lower %Rank values indicate more favorable predicted ranking. This analysis reflects prediction-based assessment rather than direct experimental validation of T cell activation or clinical immunogenicity. Abbreviations: NES, Normal-Epitope Sources; PBMC, peripheral blood mononuclear cells. Alt text: Figure 2 summarizes TAE and TSE distributions across cancer types, overlap with public immunopeptidomics databases, epitope category composition, event-class contributions, predicted binding affinities, and allele-specific immunogenicity rankings.

Mapping epitope occurrence across cancer types (Figure 2C) revealed cancer-lineage-restricted patterns, with 79.3% of TAEs/TSEs detected in a single cancer lineage. We also identified 65 conserved TSEs recurrent across more than 10 tumor types, representing potential shared antigen candidates for future pan-cancer vaccine prioritization; Supplementary Tables S4 and S5 list TSEs and TAEs identified in more than 10 cancer types. Compositional analysis established TAEs as the dominant class (83.8% of total tumor epitopes), with maximal enrichment in melanoma, leukemia, meningioma, lymphoma, and breast cancer, where TAE abundance was consistent with cognate gene transcription, and top genes were highlighted by expression/epitope density (Figure 2D).

Integrated proteogenomic analysis further identified candidate tumor-associated antigens per cancer type (Supplementary Table S6), including *PMEL* in melanoma [42], *CSPG4* [43,44], *CD74* in B-cell lymphomas [45], and *HSP90A* in breast cancer [46]. In parallel, recurrent TSE-associated events supported by available TCGA genomic or transcriptomic recurrence information and immunopeptidomics detection were prioritized across representative cancer types (Supplementary Figure S4). TSEs predominantly originated from non-canonical transcriptional or translational sources, with 3′ overlapping ORFs generating 16,403 epitopes, processed pseudogenes contributing 11,483 epitopes, and 5′ upstream ORFs yielding 10,436 ligands (Figure 2D). We also compiled the top 50 epitopes for each cancer type ranked by %Rank to facilitate cancer-type-specific candidate prioritization (Supplementary Table S7). Binding-affinity analysis showed that both classes contained predicted binders (affinity < 500 nM), with TSEs showing stronger predicted binding affinity than TAEs for HLA-A and HLA-B alleles (42.63 nM vs. 61.18 nM; Figure 2E). Similar trends were observed for HLA-DP (Supplementary Figure S5). Predicted immunogenicity assessment across high-frequency HLA allotypes revealed TSE-dominant patterns in HLA-A*01:01 and HLA-B*07:02 contexts and TAE-dominant patterns in HLA-A*02:01, HLA-B*24:02, and HLA-B*27:05 contexts (Figure 2F), potentially reflecting allele-dependent peptide preferences such as conserved HLA-B*27:05 P2-Arg anchoring. These analyses show how NeoOmni Atlas can be used to compare TAE/TSE distributions, non-canonical source classes, and predicted HLA-associated properties across cancer types.

### Usage of NeoOmni Atlas

NeoOmni Atlas provides interactive modules for searching, browsing, downloading, and annotating immunopeptidomics-supported epitopes (http://modinfor.com/NeoOmni-Atlas/). The search page offers simple, advanced, peptide, HLA, event, and PTM search modes (Supplementary Figure S6), allowing both broad database exploration and targeted lookup of individual peptides, alleles, source events, or modifications. The platform also provides browsing by cancer type, HLA allele, PTM, and tissue type (**Figure 3**A). Result tables display peptide sequence, associated HLA alleles, epitope type, sample source, modification type, event classification, and event information (Figure 3B–C). Users can inspect annotated MS/MS spectra, peptide-HLA-sample/cancer networks, and HLA binding details, including binding level, affinity, ranking score, and sample/dataset information (Figure 3D–F). These features support multidimensional exploration, download, and prioritization of candidate tumor (neo)antigens while allowing users to connect peptide sequences with source events, sample context, predicted HLA presentation, and MS/MS evidence. The downloadable tables support external reanalysis, candidate filtering, and integration with independent genomic or immunological datasets.

**Figure 3.**
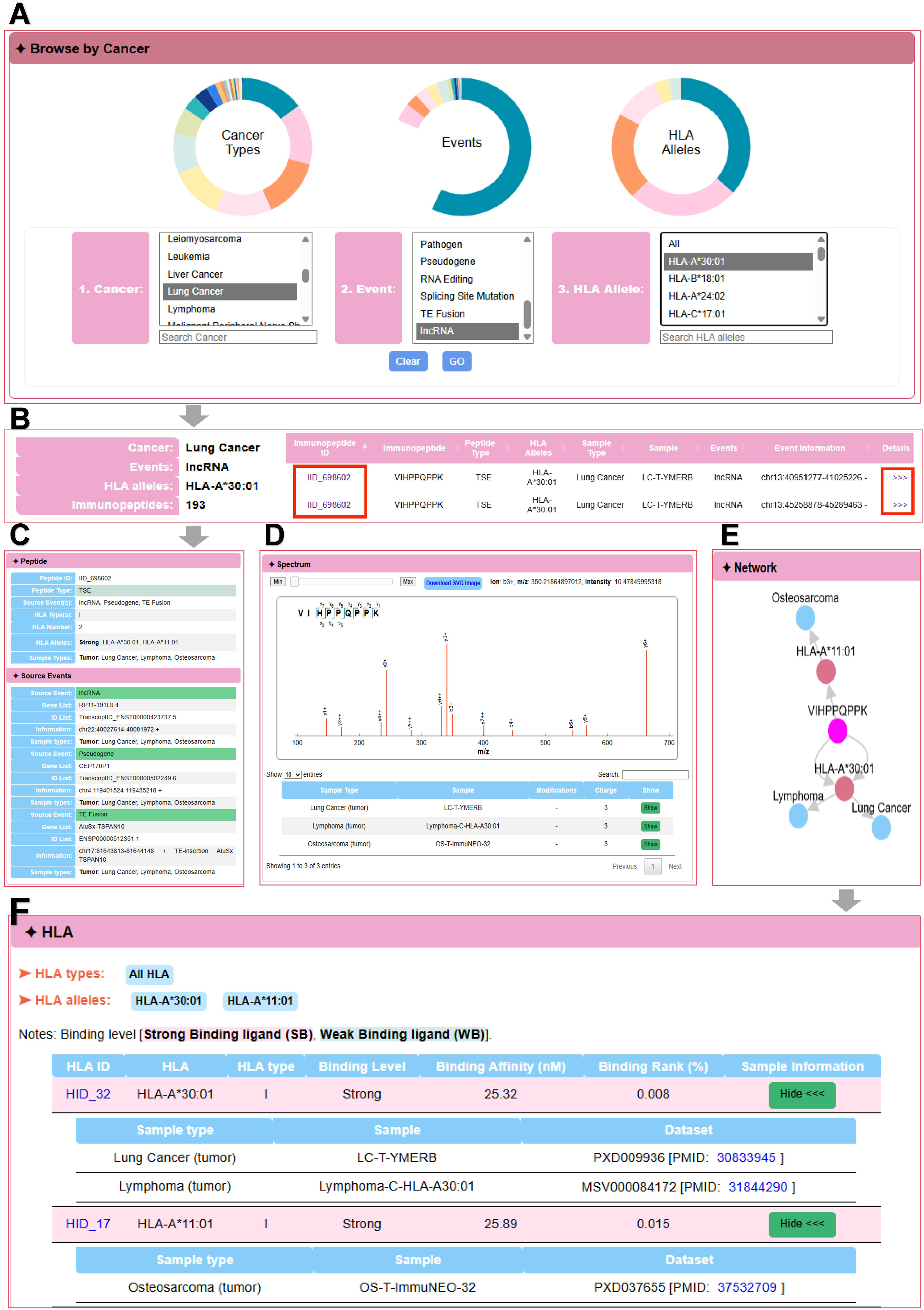
User interface of NeoOmni Atlas. NeoOmni Atlas is freely accessible at http://modinfor.com/NeoOmni-Atlas/ and provides web-based modules for browsing, searching, visualizing, and downloading immunopeptidomics-supported epitope annotations. (A) Browsing interface for exploring immunopeptides by cancer type, event type, HLA allele, post-translational modification, and tissue type. (B) Representative result table showing immunopeptides filtered by selected cancer type, event category, and HLA allele. (C) Peptide-level information page displaying key annotations, including peptide sequence, epitope type, source gene or event, sample information, cancer/tissue type, and dataset source. (D) Annotated MS/MS spectrum of the selected peptide, supporting mass spectrometry-based peptide identification. (E) Network visualization showing relationships among the peptide, predicted HLA binder, and the samples or cancer types in which the peptide was detected. (F) HLA binding details, including predicted binding level, binding affinity or ranking score, HLA allele, and sample/dataset information. Abbreviations: MS/MS, tandem mass spectrometry; PTM, post-translational modification. Alt text: Figure 3 displays the NeoOmni Atlas web interface, including browsing filters, peptide result tables, peptide annotation pages, an annotated MS/MS spectrum, a peptide-HLA-sample network, and HLA binding information.

## Discussion

NeoOmni Atlas is a large-scale immunopeptidomics-supported pan-cancer antigen resource integrating canonical peptides and 18 non-canonical/additional antigen source classes across 26 cancer types. It contains 759,386 unique HLA-presented epitopes, including 316,126 TAEs and 61,231 TSEs, and broadens dataset, cancer-type, HLA-allele, and event-annotation coverage relative to existing resources such as HLA Ligand Atlas, SysteMHC Atlas, and caAtlas [22–24]. By incorporating genomic variants, transcriptional dysregulation, RNA-level events, pathogen-derived sequences, and non-canonical translation products, NeoOmni Atlas captures antigen sources that are underrepresented in previous immunopeptidomics databases. The 65 conserved TSEs recurrent across more than 10 tumor types represent potential shared antigen candidates for further evaluation, while lineage-restricted TAEs and TSEs provide cancer-type-specific candidate pools. Together, these results support the value of integrated immunopeptidomics reanalysis for transforming heterogeneous public MS datasets into a structured, queryable antigen resource.

TAE and TSE annotations should be interpreted within the operational framework used here. TAEs are canonical tumor-derived peptides not detected in the compiled normal/benign peptide reference set, whereas TSEs are tumor-type-specific and tissue-context-dependent non-canonical peptides filtered against corresponding normal/benign references and the complete TAE peptide set. These categories support discovery-level antigen prioritization and should not be interpreted as proof of universal absence from all normal tissues or as experimental validation of clinical tumor specificity or safety. Additional comparison with the HLA Ligand Atlas normal/benign reference is provided in Supplementary Figure S7. Candidate peptides intended for therapeutic development require additional normal immunopeptidome screening, expression-level assessment, and functional safety validation.

NeoOmni Atlas also provides prioritization strategies for candidate source genes and events. For canonical TAE source genes, the composite score integrates immunopeptidomic evidence and transcriptomic abundance, highlighting genes with frequent HLA presentation and tumor-associated expression. Several prioritized genes correspond to reported tumor-associated targets, including *PMEL, CSPG4, CD74*, and *HSP90A*, supporting the ability of the atlas to rediscover known targets while nominating additional candidates. Because this score is gene-expression-based, it is primarily applicable to canonical TAE source genes; for TSE-associated events, TCGA recurrence information and MS-based detection were used as complementary event-level prioritization metrics. This design allows known antigen rediscovery and prioritization of additional candidate sources while avoiding overinterpretation of heterogeneous non-canonical event annotations.

Several limitations remain. Immunopeptidomics sensitivity depends on sample quantity, instrument performance, sample preparation, HLA enrichment efficiency, and processing pipelines, so low-abundance epitopes may be missed. HLA assignment relies on genotype-informed prediction and deconvolution and should be viewed as predicted association rather than definitive experimental restriction. In addition, the 18 non-canonical/additional antigen source classes were derived from multiple public resources rather than fully matched paired sequencing data from the same immunopeptidomics samples, and cancer-type coverage remains biased toward public data availability. Moreover, predicted binding or presentation potential should not be equated with experimentally validated T cell recognition or clinical immunogenicity. Additional matched sequencing data, mono-allelic immunopeptidomics, rare-cancer datasets, and normal/benign references will improve future updates and help distinguish broadly recurrent candidates from context-dependent or potentially cross-reactive peptides. Overall, NeoOmni Atlas provides a searchable and downloadable resource for MS-supported tumor (neo)antigen candidate prioritization and future immunotherapy studies. Future versions will benefit from additional normal/benign references, quantitative presentation information, longitudinal sampling, and experimentally validated peptide-HLA interactions, which will further refine candidate prioritization and safety assessment.

## Ethical statement

This study used publicly available and previously published datasets and did not involve new human participants, human specimens, or animal experiments.

## Data availability

The data are freely accessible and downloadable from the NeoOmni Atlas platform (http://modinfor.com/NeoOmni-Atlas/). The curated datasets can be browsed, searched, and downloaded from the platform through the Download page.

## Code availability

The corresponding author can provide the scripts used for data processing and analyses in response to a reasonable request.

## CRediT author statement

**Tao Zheng**: Conceptualization, Data curation, Methodology, Writing – original draft. **Zhi Ran**: Conceptualization, Data curation, Methodology, Writing – original draft. **Jing Zeng**: Formal analysis, Data curation. **Tao Wang**: Formal analysis, Methodology. **Shiyang Fu**: Validation, Data curation. **Lihua Cai**: Validation, Data curation. **Dinghong Wu**: Software, Visualization. **Jiaxiang Ding**: Software, Visualization. **Tang Liu**: Funding acquisition. **Zhihong Li**: Formal analysis, Validation. **Kai Yuan**: Supervision, Project administration, Writing – review & editing. **Shaofeng Lin**: Supervision, Software, Visualization, Writing – review & editing. **Haodong Xu**: Supervision, Project administration, Funding acquisition, Writing – review & editing. The final version was read and approved by every author.

## Acknowledgments

This study was supported by the Excellent Youth Foundation of Hunan Scientific Committee (Grant No. 2024JJ2084), National Natural Science Foundation of China (Grant No. 32300528, 82472448, 92153301, 32170821, 32370972, 32300520, and 82402824), the Scientific Research Fund of Hunan Provincial Education Department (Grant No. 23B0023), the Science and Technology Innovation Program of Hunan Province (Grant No. 2023RC3080), Central South University Research Programme of Advanced Interdisciplinary Studies (Grant No. 2023QYJC026), the Scientific Research Foundation for Advanced Talents of Fujian Medical University (Grant No. XRCZX2022015), the National Key Research and Development Program of China (Grant No. 2021YFC2701200), the Department of Science & Technology of Hunan Province (Grant No. 2023RC1028, 2023SK2091, and 2021JJ10054), and the Graduate Innovation Project of Central South University (Grant No. 2024XQLH056 and 2025ZZTS0306). The funders had no involvement in designing the study, collecting or analyzing data, preparing the manuscript, or deciding on publication. We thank the investigators who made their immunopeptidomics datasets publicly accessible, enabling this integrated analysis.

## Supplementary material

Supplementary material includes Supplementary Figures S1–S7 and Supplementary Tables S1–S7. These materials are provided as separate supplementary files and are cited in the main manuscript where applicable.

**Supplementary Figure S1. Workflow for event-specific template construction.**

Schematic workflow illustrating the construction of non-canonical event-specific candidate amino-acid sequence templates for the customized search space. The event categories shown in the schematic represent grouped examples of the 18 non-canonical/additional antigen source classes summarized in Supplementary Table S2. Reference genome/transcriptome annotations, reference proteome information, and tumor-derived variant or event annotations were used as input resources. Event-specific nucleotide templates were constructed according to event type, including ORF-specific sequence extraction, mutant-centered sequence window construction, junction- or fusion-spanning template construction, and extraction of non-coding or intronic source sequences. The resulting templates were translated in a strand-aware and frame-aware manner, with reverse-complement frames considered where applicable, to generate non-canonical event-specific candidate amino-acid sequence templates.

**Supplementary Figure S2. Absolute and search-space-normalized peptide yield across non-canonical event classes.**

(A) Number of identified unique peptides across the 18 non-canonical event classes. (B) Search-space-normalized peptide yield, calculated as the number of identified unique peptides per one million candidate 8 – 25 amino-acid windows generated from each event class. This normalization accounts for differences in event-class-specific search-space size and provides a complementary view of peptide recovery beyond absolute peptide counts.

**Supplementary Figure S3. Comparison of the NeoOmni Atlas with other public immunopeptidomics resources.**

(A) Overlap of datasets among the NeoOmni Atlas, IEAtlas, SysteMHC Atlas, and caAtlas. (B) Overlap of cancer types among the four resources. (C) Overlap of HLA class I alleles among the NeoOmni Atlas, HLA Ligand Atlas, SysteMHC Atlas, and caAtlas. (D) Overlap of HLA class II alleles among the four resources.

**Supplementary Figure S4. Tumor-restricted TSE-associated events identified by integration of TCGA recurrence information and immunopeptidomics detection.**

Recurrent TSE-associated events were prioritized across representative cancer types by integrating available TCGA genomic or transcriptomic recurrence information with immunopeptidomics detection. The figure summarizes prioritized events supported by both tumor recurrence information and detected TSEs. Abbreviations: TCGA, The Cancer Genome Atlas.

**Supplementary Figure S5. Binding affinity comparison of TAEs and TSEs to HLA-DP, HLA-DQ, and HLA-DR alleles.**

Consistent with the HLA class I analysis shown in Figure 2E, TSEs showed a similar trend toward stronger predicted binding affinity than TAEs for HLA-DP.

**Supplementary Figure S6. Search mode of the NeoOmni Atlas.**

(A) Simple Search: Enter a peptide sequence to retrieve detailed information, including peptide type and associated mutation events. (B) Event Search: Search for specific mutation events to explore detailed characteristics and contextual information. (C) HLA Search: Query the number and details of peptides bound to specific HLA alleles. (D) Advanced Search: Perform precise searches by combining peptide sequences and HLA alleles for tailored results. (E) PTM Search: Retrieve sample information related to PTMs of proteins, highlighting modification features. (F) Peptide Search: Access sample information associated with a specific peptide, revealing its distribution and characteristics across samples.

**Supplementary Figure S7. Area-proportional two-set Euler diagrams show the peptide sequence-level overlap between NeoOmni antigen candidates and the HLA Ligand Atlas normal/benign reference set.**

(A) Comparison of NeoOmni TAEs with the HLA Ligand Atlas reference set. (B) Comparison of NeoOmni TSEs with the HLA Ligand Atlas reference set. Numbers indicate unique peptide sequence counts in each region. Percentages indicate the proportion of NeoOmni TAE or TSE peptides not detected in the HLA Ligand Atlas reference set.

**Supplementary Table S1. Metadata of immunopeptidomics datasets included in NeoOmni Atlas.**

**Supplementary Table S2. Summary of 18 classes of non-canonical neoantigen-generating events included in this study.**

**Supplementary Table S3. Modifications selected in our database.**

**Supplementary Table S4. TSEs detected across more than 10 cancer types.**

**Supplementary Table S5. TAEs detected across more than 10 cancer types.**

**Supplementary Table S6. Actionable tumor antigens identified across cancer types.**

**Supplementary Table S7. Top 50 predicted epitopes per cancer type.**

