## Supplementary figures and images for "NeoOmni Atlas: an integrated database for pan-cancer neoantigen derived from proteogenomics analysis"

### Supplementary Figure S1

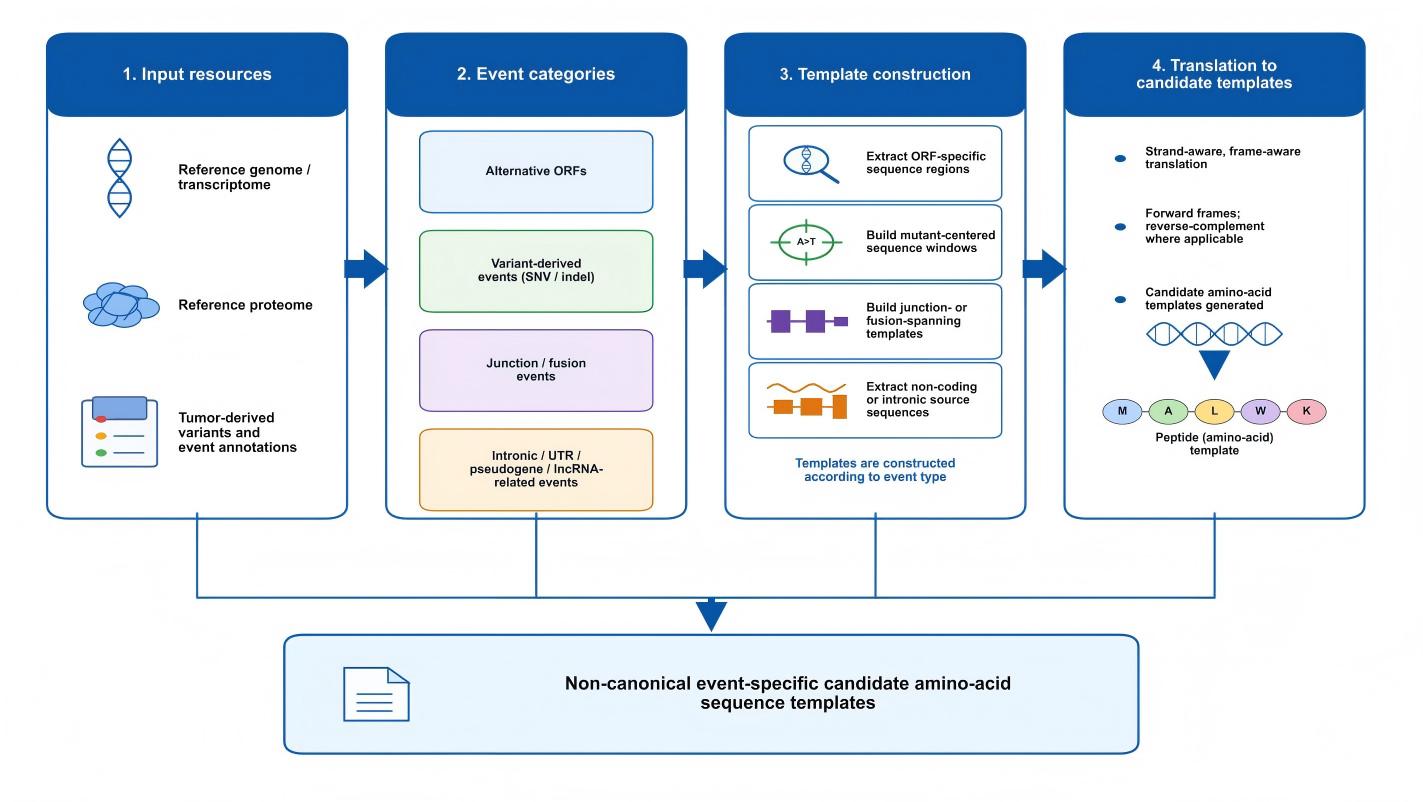

### Supplementary Figure S2

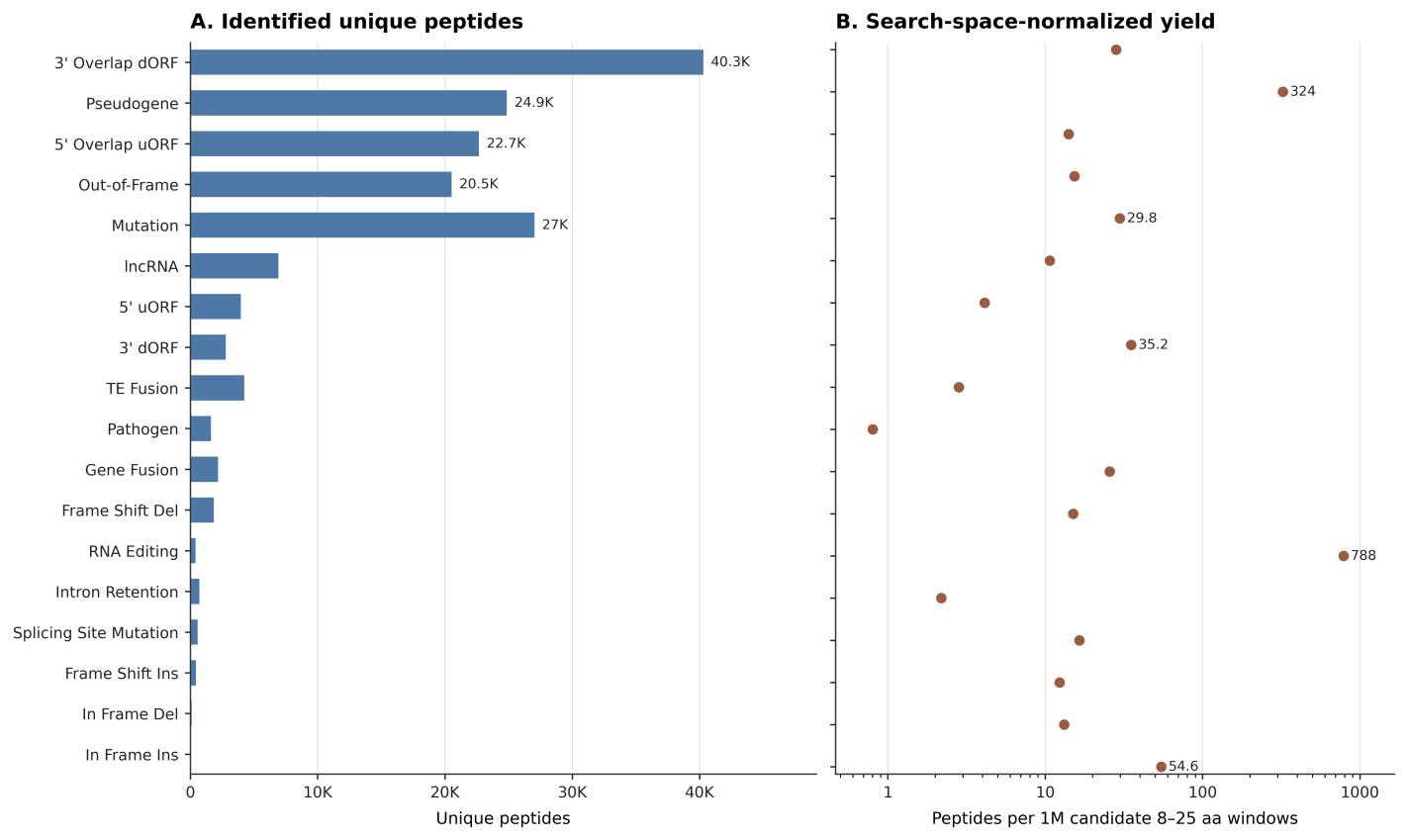

### Supplementary Figure S3

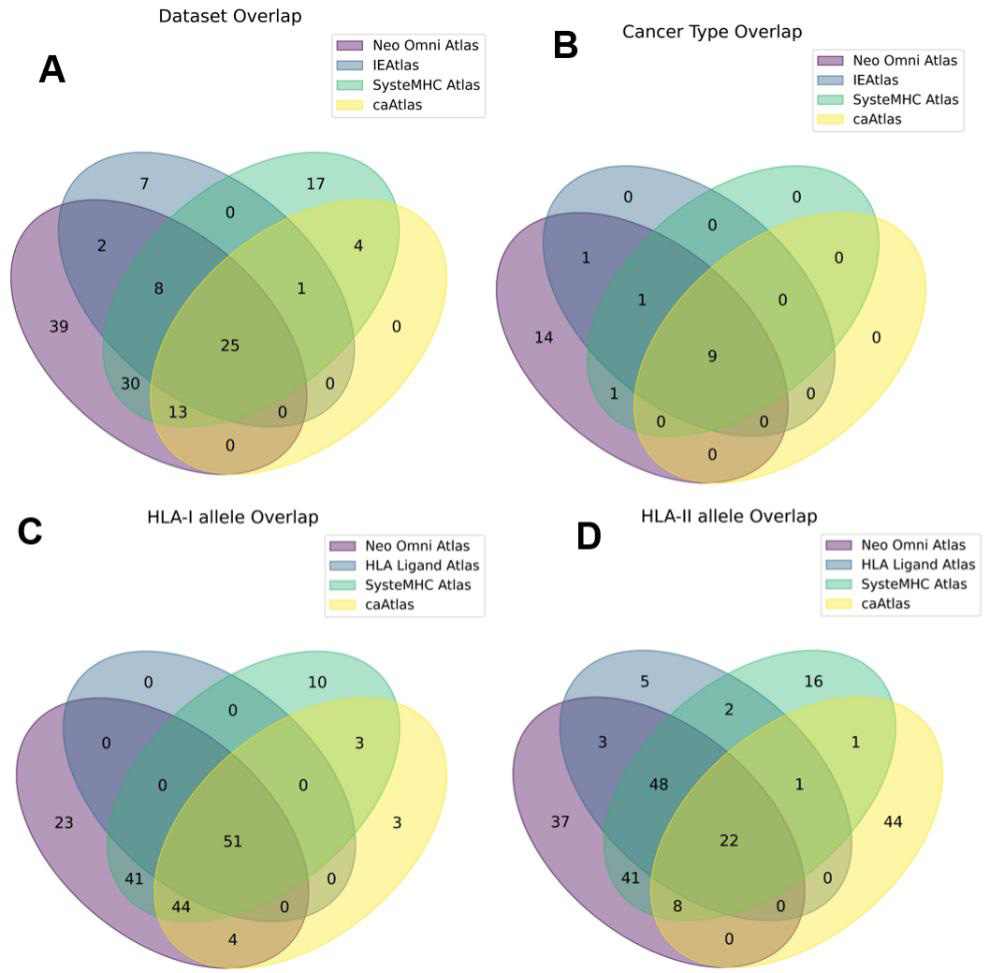

### Supplementary Figure S4

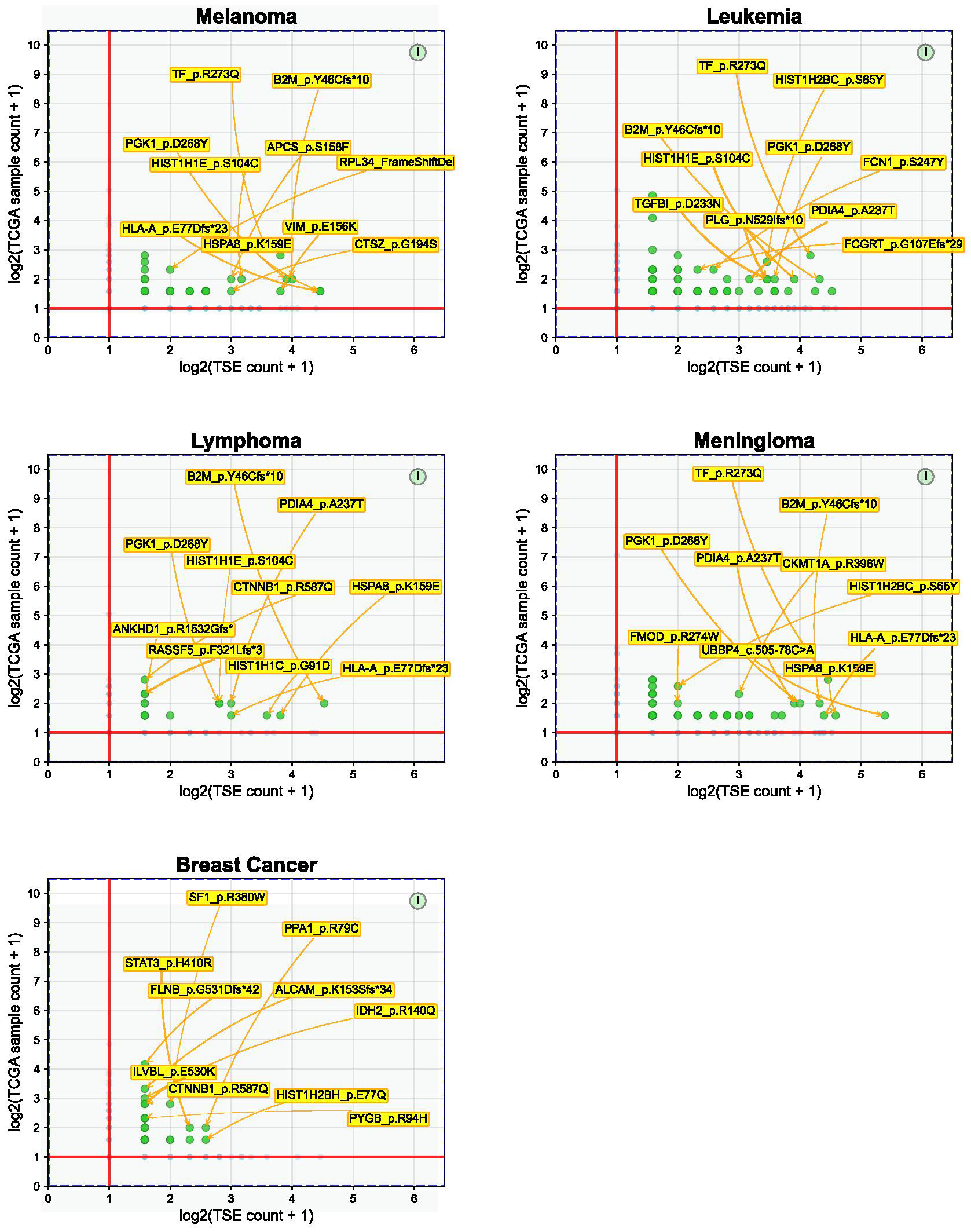

### Supplementary Figure S5

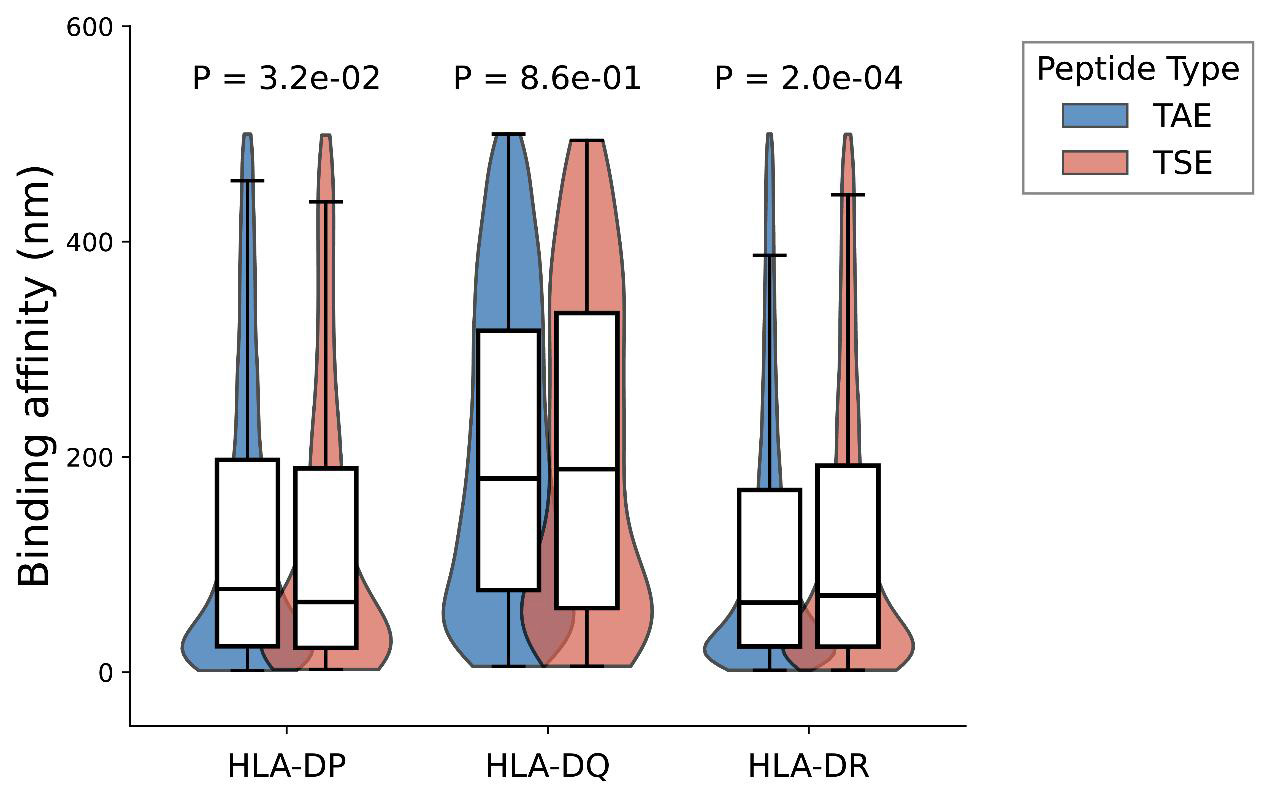

### Supplementary Figure S6

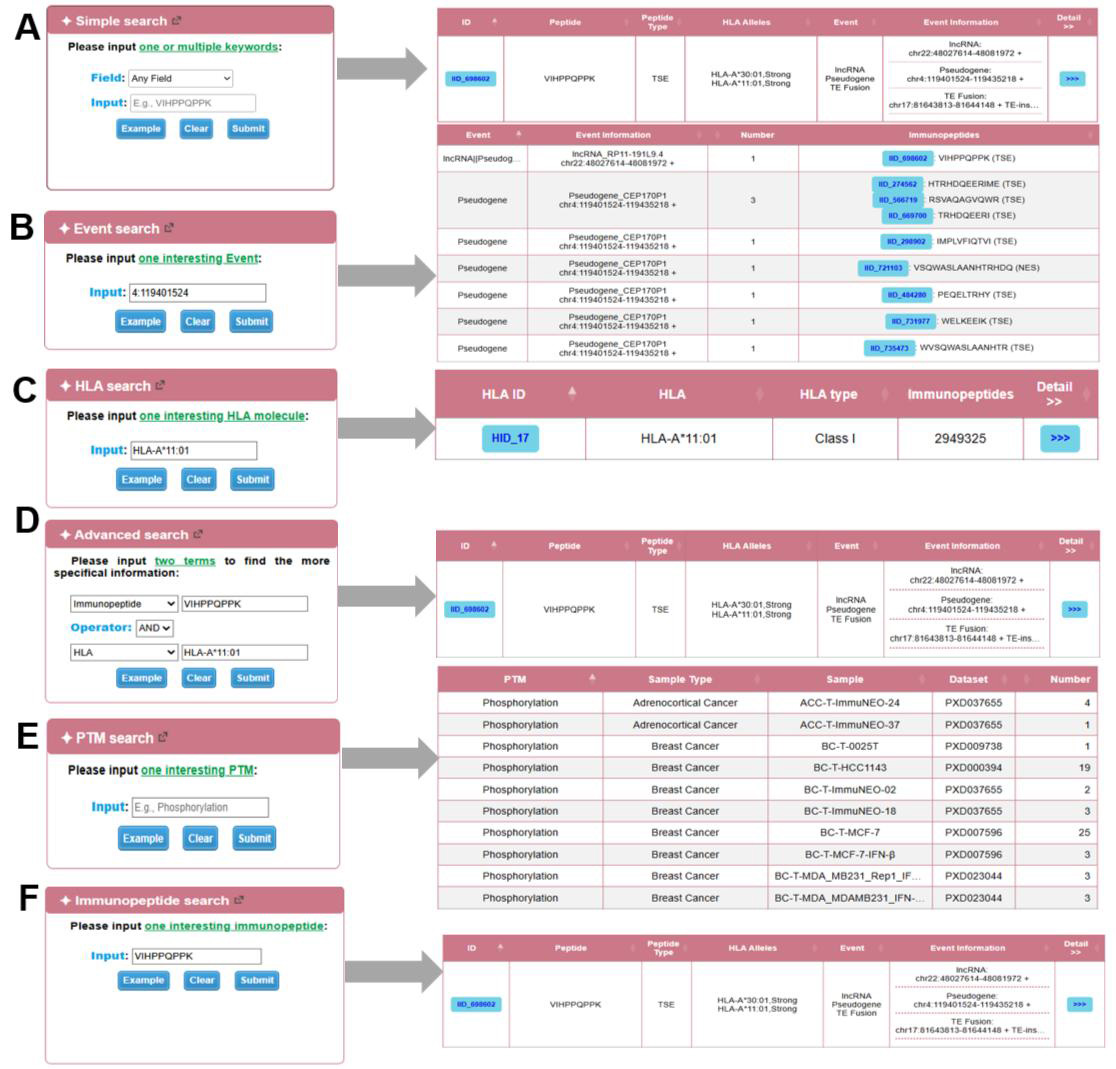

### Supplementary Figure S7

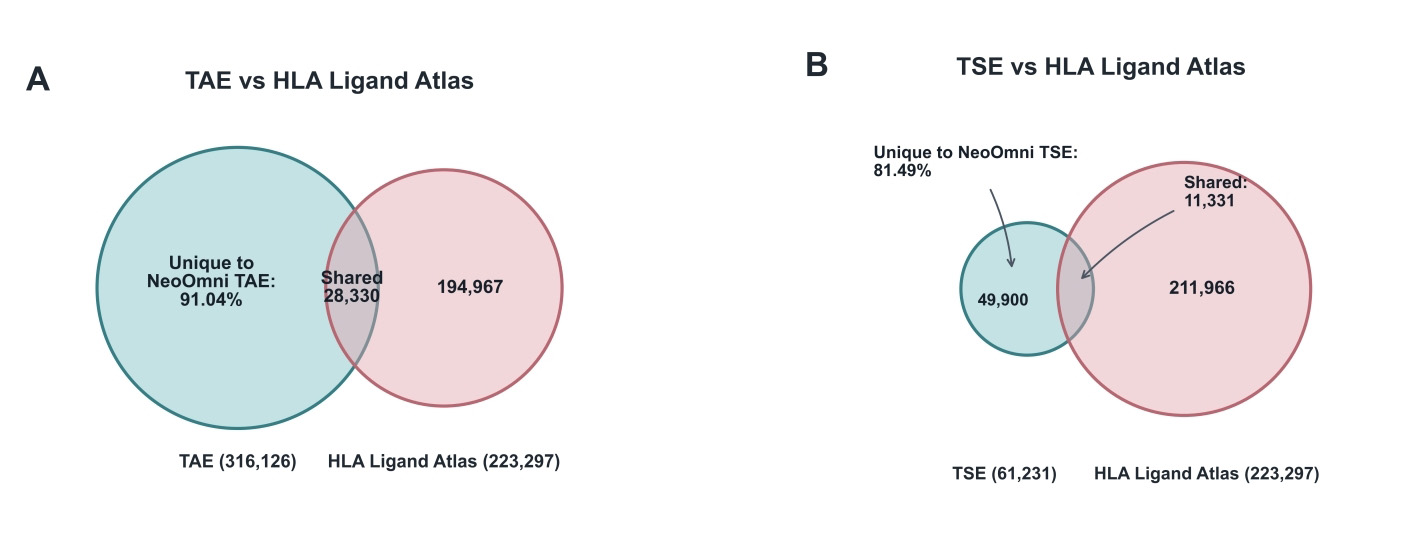
